# The multiplier benefits of integrating non-sewered and sewered wastewater treatment and sanitation processes

**DOI:** 10.1101/2025.07.22.666153

**Authors:** Doulaye Kone, Liron Friedman, Kartik Chandran

**Author notes:** Correspondent footnote:* Kartik Chandran, Department of Earth and Environmental Engineering, Columbia University, 500 West 120th Street, New York, NY 10027., URL: www.columbia.edu/~kc2288.

## Abstract

This study showcases the beneficial integration of non-sewered sanitation systems (NSSS) with sewered wastewater treatment plants (WWTPs). Treating increasing fractions of influent wastewater loads via six different types of NSSS offered correspondingly increasing savings in operating energy costs at five WWTPs, employing a broad range of typically employed treatment processes. Two NSSS that treat both greywater and blackwater (gb-HRT) and blackwater alone (b-HRT) yielded the highest savings in annual operating energy costs across most WWTPs. Distinctly, NSSS involving urine-separation with and without internal recirculation promoted energy-positive operations, by enhancing anaerobic digestion in selected WWTPs. At the highest NSSS coverage tested (treating 50% of the influent sewage), savings in annual sewered WWTPs operating energy costs ranged from $76k to $800k and increased further to the range $301k to $1.1M annually with process optimization. Therefore, integration of NSSS with sewered WWTPs can improve overall treatment efficiency, while facilitating resilient sanitation practices.

## Introduction

Appropriate wastewater management plays a pivotal role in providing universal access to safe water and sanitation and in turn supporting human livelihoods, healthy ecosystems and promoting sustainable development (Prüss et al. 2002, Narain 2012, WHO et al. 2018, Kone 2023). Sewered wastewater treatment plants (WWTPs) have been successfully applied worldwide to protect environmental quality enhance human health (Lofrano et al. 2010). Nevertheless, sewered WWTPs face multiple challenges stemming from aging treatment processes, stringent regulations on effluent quality, and unpredictable population growth in the associated sewer sheds (Kone 2023). Traditional approaches to handing such increasingly common challenges include expensive WWTP expansions, retro-fits, rehabilitating or installing sewer networks afresh. Such approaches require substantial capital expenditure and could disrupt day-to-day activities of the impacted communities (Duque et al. 2024a). Alternatively, deployment of non-sewered sanitation systems (NSSS) to address existing or anticipated treatment complexities offers a more tailored solution that could have a lower initial capital outlay, permit more timely implementation installed and impose relatively less disruptions(Kone 2023). In addition, implementation of NSSS in dense urban settings offers closed-loop local reuse opportunities such as in landscape irrigation, local urban farming, and toilet flushing, among others ^(Mitra et al. 2022, Christou et al. 2024)^. Recent studies have explored the benefits of integrating NSSS to complement, substitute or enhance conventional sewered WWTP infrastructure ^(Strande et al. 2023, Garrido-Baserba et al. 2024)^. These studies have examined various facets including the cost of such integrated designs, as well as the associated influences of the topography on the cost of sewer installations and maintenance (Hesarkazzazi et al. 2022), and prospect of WWTP optimization (Zhang et al. 2023, Duque et al. 2024b). However, the quantitative impacts on integrating NSSS and centralized WWTPs in terms of process performance, treatment efficiency, associated with modifications in the influent water quality and wastewater flowrate have received little attention. Specifically, offsets in influent flows and loads as effected by upstream implementation of NSSS could directly impact downstream centralized WWTP process performance (Grady et al. 1999, Tchobanoglous et al. 2003). Furthermore, changes to influent composition and loads achieved through upstream NSSS implementation could also offer opportunities to extract more efficiency from centralized-WWTP operation resulting in improved WWTP performance with lower operating cost. Such optimization offers opportunities for utilities to reinforce the resilience of the wastewater infrastructure, especially if adoption of NSSS can help increase WWTP capacity for accommodating drainage water and minimizing the impact of flood or drought on service provision (Kone 2023).

As such different NSSS have been developed. In 2011, the Water, Sanitation & Hygiene program of the Bill and Melinda Gates foundation launched the Reinvent the Toilet Challenge that has since revolutionized the development of new **R**einvented **T**oilet (**RT**) technologies that can safely and effectively manage human waste. Implementation of such NSSS and others is aimed at enhancing access to sanitation globally and facilitate progress towards attaining targets specified in the United Nations Sustainable Development Goals.

In this study we present a novel approach using mechanistic process modeling to evaluate the effect of integrating various NSSS with a range of traditionally employed centralized WWTP configurations. Specifically, this study focuses on quantifying the potential impacts on the centralized facility in terms of (1) process performance, (2) energy savings, and (3) annual operational costs. Opportunities for process optimization following NSSS integration in terms of potential cost savings are also examined.

As part of this study, we hypothesized that the beneficial impact of different NSSS when integrated upstream of WWTPs would be a function of the specific compositional changes effected by the NSSS and their degree of implementation. For instance, reduction of influent nutrient (nitrogen and phosphorus) loads through NSSS implementation would be preferred in conjunction with WWTPs with stringent nutrient discharge targets. We further hypothesized that the increase in efficiency of optimized WWTP process would correspond to the degree of reduction in influent wastewater loads, in-turn related to the extent of NSSS implementation.

These hypotheses were tested by replicating five real existing sewered-WWTPs in South Africa, Asia and North America (Table I), using a process simulation tool (BioWin 6.2, EnviroSim Associates, ON, CA) and mechanistically determining the impact of different implementation levels of various NSS technologies (Tables II and III) on downstream WWTP performance (workflow presented in Figure 1).

**Figure 1.**
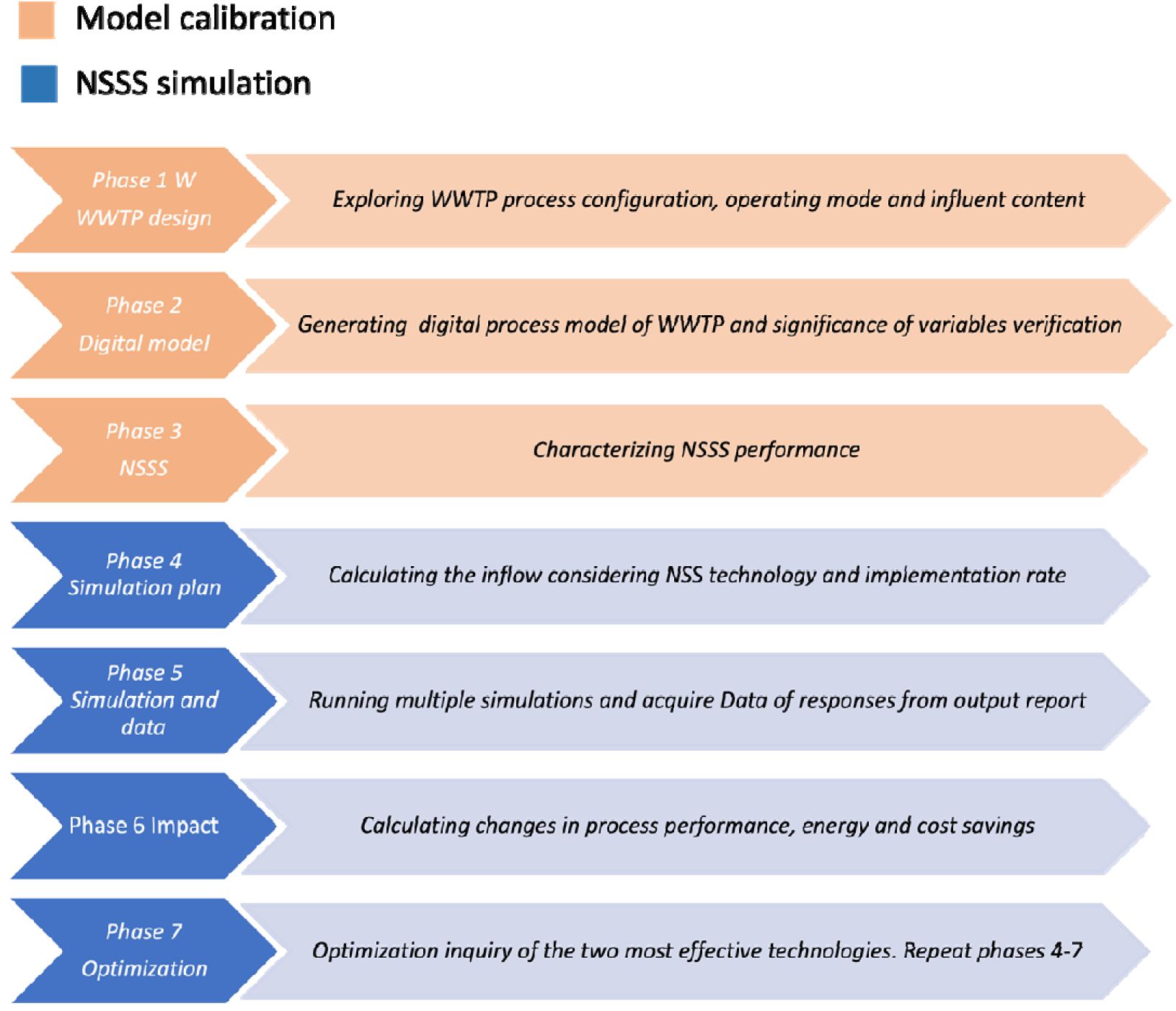
Workflow of modeling-based evaluation followed during this work. Phases colored in orange refer to calibration of the model and phases colored in blue refer to the simulation.

**Table I.** Principal features of the centralized WWTPs used in this study.

| WWTP | Location | Liquid-treatment process | Biosolids management process | Average influent characteristics |  |  |  | Population served |
| --- | --- | --- | --- | --- | --- | --- | --- | --- |
|  |  |  |  | Flow rate<br>(Q, ML/d) | Total COD<br>(mg/L) | Total Kjeldahl N (TKN, mg-N/L) | Total phosphorus<br>(mg-P/L) |  |
| **BNR1** | Beijing, China | Four-stage BNR | Not applicable | 60 | 233 | 45 | 4 | 190,433 |
| **BNR2** | Durban, South Africa | Modified Ludzack-Ettinger | Anaerobic digestion of primary sludge | 44.3 | 842 | 58.9 | 11 | 328,178 |
| **MBR** | Greater Washington, DC, USA | BNR coupled with MBR | Anaerobic digestion of thickened primary and secondary sludge | 30.3 | 630 | 60 | 8 | 195,319 |
| **WR1** | California, USA | Integrated, trickling filter, nitrification and BNR | Anaerobic digestion of thickened primary and secondary sludge | 454 | 488 | 56 | 6.5 | 2,718,127 |
| **WR2** | California, USA | Integrated trickling filter and HPOAS | Anaerobic digestion of thickened primary and secondary sludge | 223.3 | 488 | 56 | 6.5 | 1,336,323 |

**Table II.**
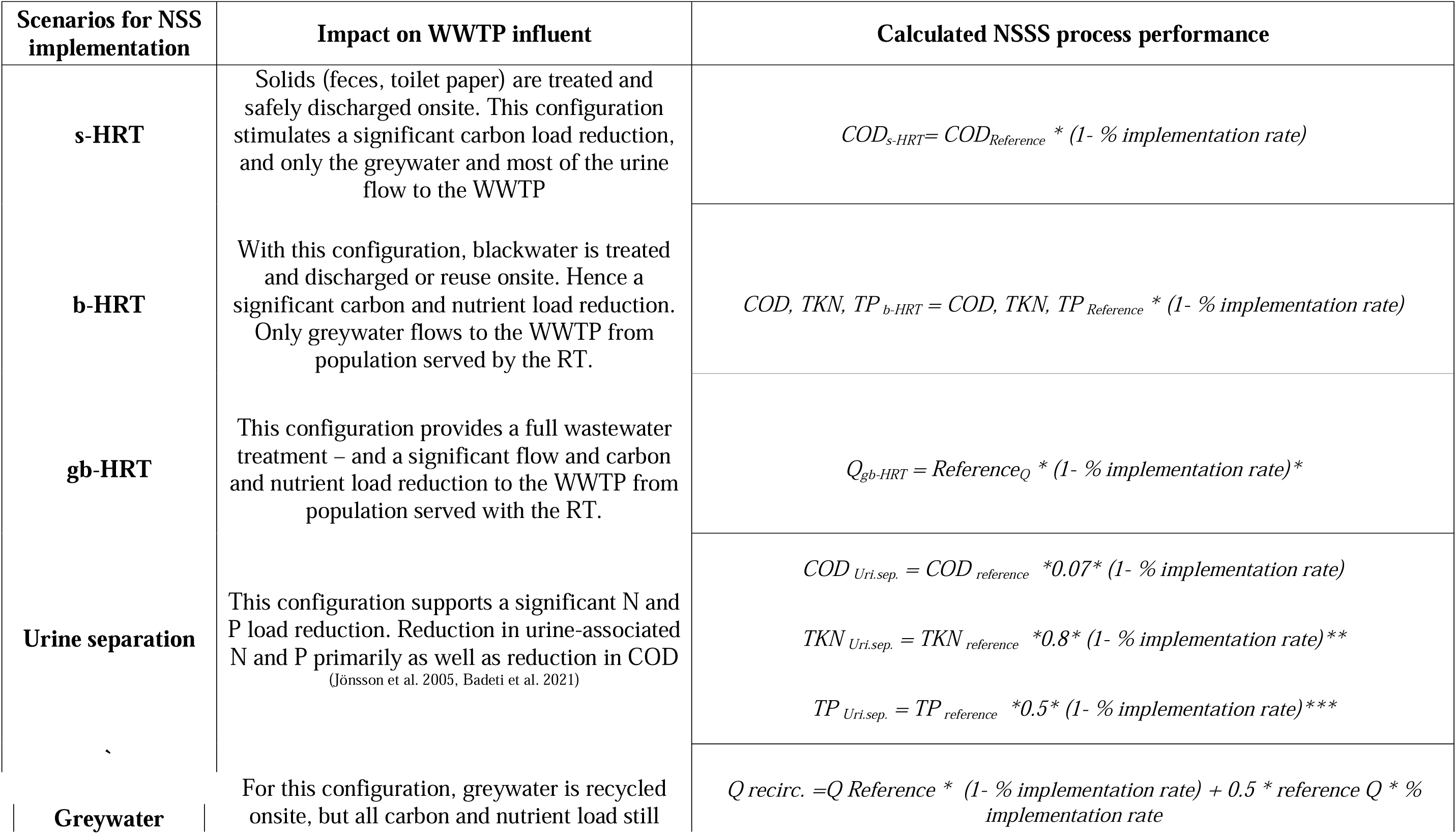
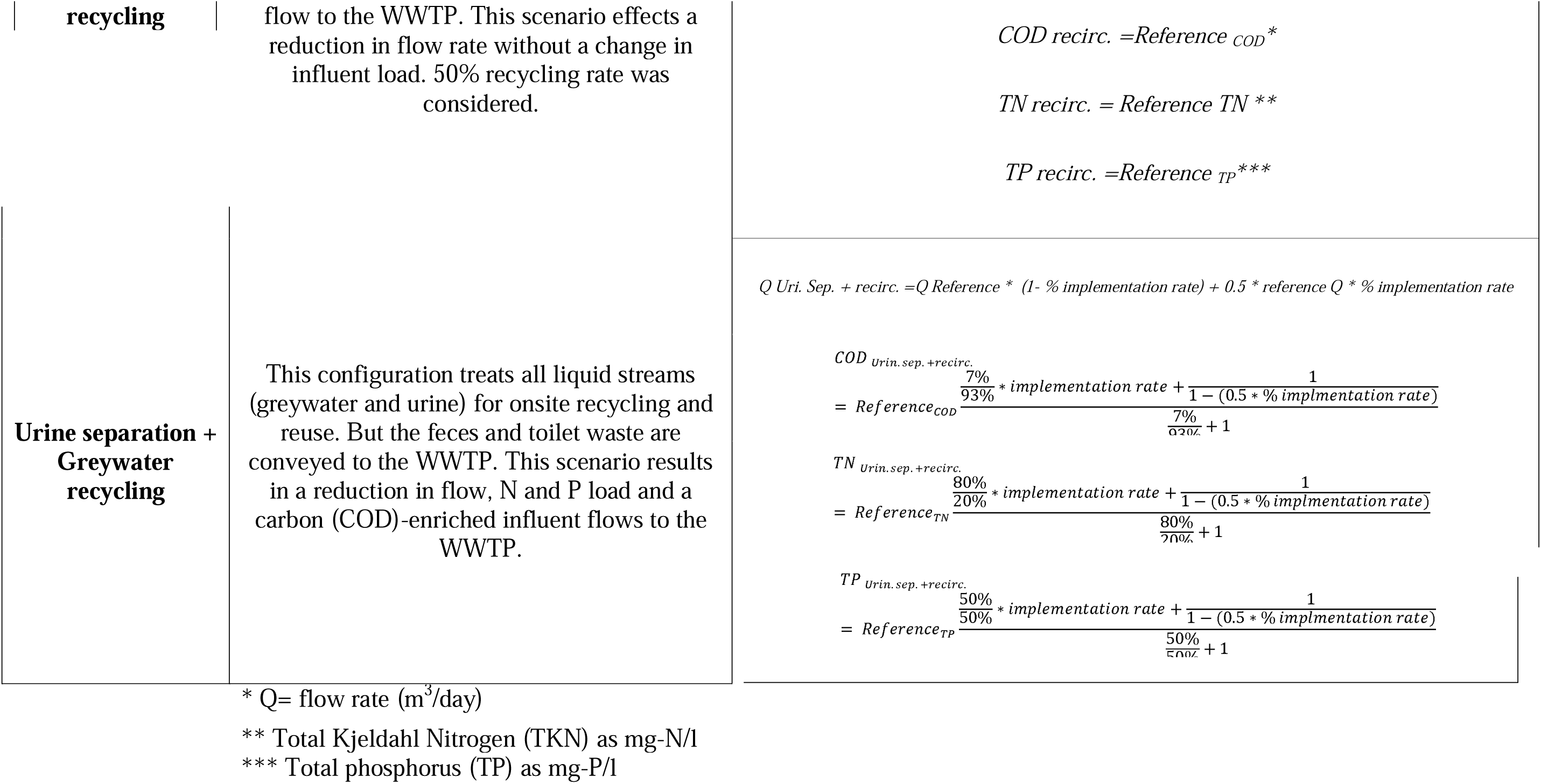
NSS scenarios evaluated.

**Table III.** Urine characteristics used for simulations that include urine separation.

| Urine characteristic | Percentage of domestic<br>wastewater (%) | Reference |
| --- | --- | --- |
| N (mg/L) | 80 | (Wuang et al. 2013) |
| P (mg/L) | 50 | (Wuang et al. 2013) |
| COD (mg/L) | 7 | (Badeti et al. 2021) |
| TSS (mg/L) | 2 | (Jönsson et al. 2005) |
| Flow rate (L/capita-d) | 2 | (Jönsson et al. 2005) |

## Results and Discussion

In general, most, but not all upstream NSS implementation scenarios resulted in better energy-efficiency at existing downstream sewered WWTPs owing to influent load reductions and compositional changes, for instance changing ratios of influent chemical oxygen demand to nitrogen, (COD:N, Figure 2 and Table IV). Additionally, the reduced influent loading resulting from NSSS implementation had differing impacts on the downstream process performance based on (a) the specific type of NSS system and (b) the specific configuration and discharge limits or performance requirements of the downstream system. Interestingly, our analysis also highlighted inefficiencies relating to the current operations and potential opportunities for improving the efficiency of the existing WWTPs. Next, we describe the specific impacts of the various NSSS on the different WWTPs by comparing changes from the simulation outputs of the integrated NSSS-centralized systems (operational cost, energy demand, process performance) to the outputs of the centralized-only scenario.

**Figure 2.**
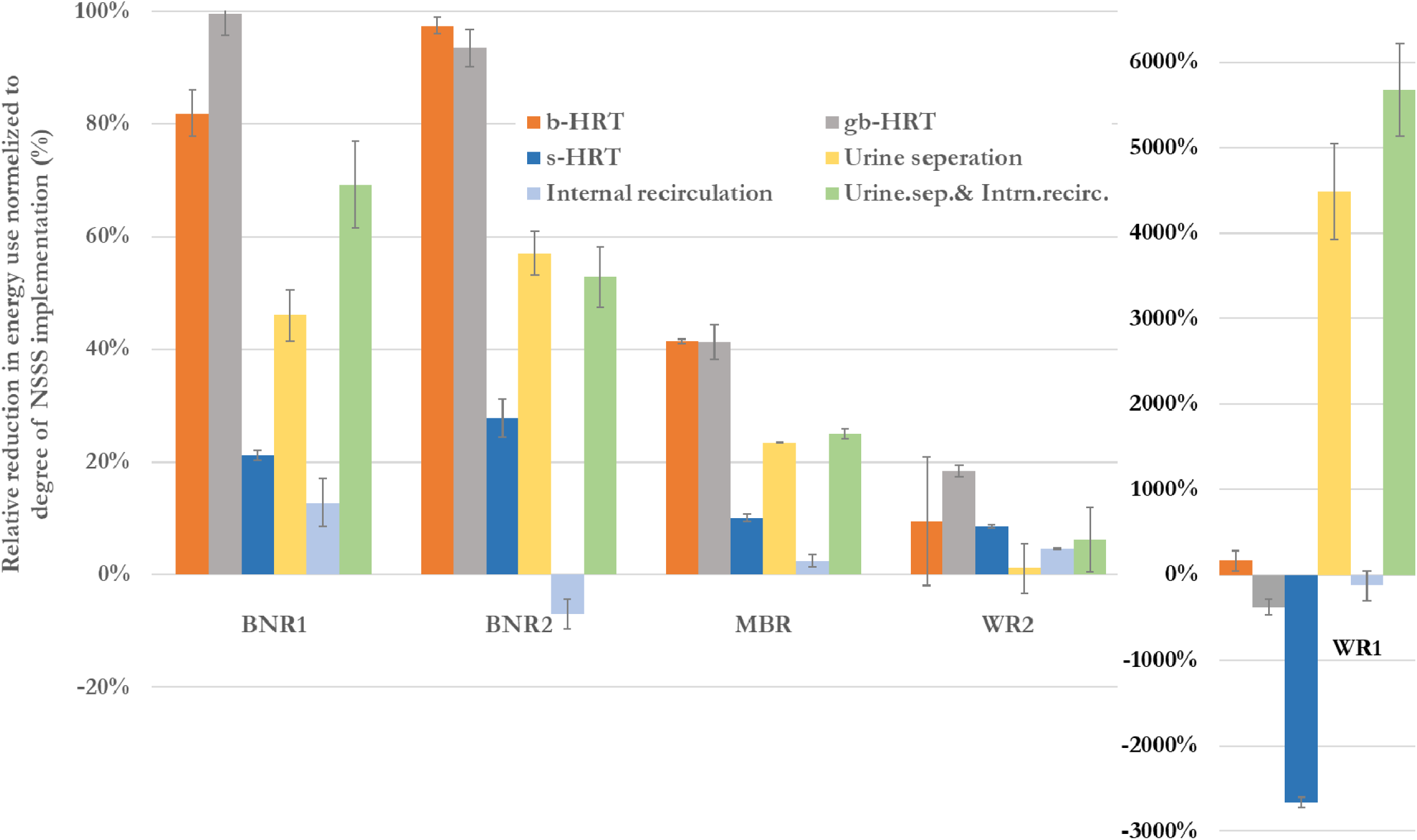
Relative savings in per capita energy costs normalized to degree of implementation (%) at the different sewered WWTPs tested as a function of different degrees of NSSS implementation <u>without optimization</u>. Negative savings indicate an increase in operating energy costs. Height of the bars represents relative savings averaged over 10%, 30% and 50% NSSS implementation. Error bars represent one standard deviation.

**Table IV.** Best intervention for energy savings for 10%, 30%, and 50% population coverage for current and optimized operations.

| Coverage | Plant | Without optimization |  |  | With optimization |  |  | Fold-benefit of optimization |
| --- | --- | --- | --- | --- | --- | --- | --- | --- |
| | | Best Intervention | Energy Savings (%) | Monetary Savings (\$) | Best Intervention | Energy Savings (%) | Monetary Savings (\$) | |
| 10% | BNR1 | gb-HRT | 10% | \$82,312.74 | gb-HRT | 36% | \$287,240.40 | 3.5x |
| | BNR2 | b-HRT | 10% | \$57,768 | b-HRT | 25% | \$138,671 | 2.4x |
| | MBR | gb-HRT | 4% | \$22,791.60 | gb-HRT | 39% | \$201,392.40 | 8.8x |
| | WR1 | US+Int | 609% | \$192,852.00 | US+Int | 2001% | \$632,647.20 | 3.3x |
| | WR2 | gb-HRT | 2% | \$21,389.04 | gb-HRT | 56% | \$561,778.80 | 26x |
| 30% | BNR1 | gb-HRT | 30% | \$236,506.68 | gb-HRT | 48% | \$379,658.40 | 1.6x |
| | BNR2 | b-HRT | 31% | \$170,499 | gb-HRT | 44% | \$240,988 | 1.6x |
| | MBR | gb-HRT | 13% | \$65,043.72 | b-HRT | 51% | \$264,201.60 | 4.2x |
| | WR1 | US+Int | 1760% | \$556,903.98 | US+Int | 2913% | \$921,201.60 | 1.7x |
| | WR2 | gb-HRT | 6% | \$55,664.10 | gb-HRT | 58% | \$577,897.20 | 10x |
| 50% | BNR1 | gb-HRT | 48% | \$381,058.02 | gb-HRT | 60% | \$479,588 | 1.3x |
| | BNR2 | gb-HRT | 47% | \$260,526 | gb-HRT | 60% | \$330,478 | 1.3x |
| | MBR | b-HRT | 20% | \$103,088.16 | gb-HRT | 59% | \$301,112 | 3.1x |
| | WR1 | US+Int | 2530% | \$800,598.78 | US | 3585% | \$1,134,495.72 | 1.5x |
| | WR2 | gb-HRT | 8% | \$76,001.22 | gb-HRT | 59% | \$590,565.42 | 7.8x |

### Impact of NSSS integration on cost savings

#### Biological nitrogen removal 1, (BNR1): Beijing, China

Of the different upstream NSS options tested for integration with the BNR1 process, the household re-invented toilet (HRT) that treats both graywater and blackwater (gb-HRT) presented the highest benefit in terms of operating cost savings due to the <u>parallel</u> reduction in both influent flow and influent concentrations (Figure 2, Table IV). The blackwater-HRT (b-HRT) presented the next best option (Figure 2). For b-HRT, the benefits were primarily by virtue of treating an influent wastewater with a lower concentration but at the same flow rate (lower load). A consequence of such an operation could be the possibility to expand the capacity of similarly configured WWTPs, but not to the extent possible with gb-HRT due to an upper bound on secondary clarification in terms of surface overflow rates. On the other hand, WWTPs downstream of b-HRT processes could also be tailored to accept internal recycle, agricultural, food- and animal-waste inputs on one hand, or, to target performance to potable reuse on the other hand. The ability of WWTPs downstream of b-HRT processes to buffer stormwater inputs (in isolated tankage) is also expected to be lower than that of WWTP downstream of gb-HRT processes due to the lower flowrate reduction with b-HRT than with gb-HRT.

At the other end of the spectrum, internal recycling of the treated water with the non-recycled water did not result in a substantial change in operating costs (Figure 2). It should be noted that the internal recycling scenario resulted in a net decrease in influent flow to the downstream WWTP without a change in load (higher influent concentrations). The lack of a substantial impact of internal recirculation essentially shows that performance measures should be invariant <u>at a given WWTP solids residence time (SRT)</u> if the influent load does not change, which aligns well with fundamental wastewater design concepts (Grady et al. 1999, Tchobanoglous et al. 2003).

The beneficial impact of the HRT selectively treating solids (sHRT) was also low (Figure 2) due to the selective removal of solids (and associated organic carbon), without a concomitant reduction in influent nutrient load (nitrogen and phosphorus). Implementation of sHRT systems is also not recommended upstream of WWTP with low effluent nutrient discharge limits due to the potential limitation in organic carbon needs for BNR associated with this option. The impacts of urine separation and urine-separation coupled with internal water recycling were intermediate (Figure 2) between those associated with gb-HRT (highest benefit) and just internal recycling (minimal impact).

The benefits to upstream urine diversion were related to a reduction in operating energy costs for BNR1, again, due to an overall influent N and P load reduction (Figure 2). However, the benefits were lower than that of gb-HRT and b-HRT systems since urine contains most but not the entirety of the nitrogen and phosphorus inputs (Table III). The combination of internal recycling and urine diversion further reduced the benefits associated with urine diversion (Figure 2).

#### Biological Nitrogen Removal 2 (BNR 2), Durban, South Africa

As with the BNR1 process, the two NSSS scenarios that resulted in the highest energy savings without optimization were for gb- and b-HRT (Figure 2, Table IV). However, the presence of anaerobic digestion at the WWTP resulted in some distinctions for the different NSSS tested. Essentially, through anaerobic digestion, a fraction of the influent organic carbon (either present in the primary sludge or in the secondary sludge) can be converted to biogas for energy recovery. Consequently, the reduction of influent particulate organic carbon through the implementation of s-HRT decreased the energy recovery possible through anaerobic digestion. Nevertheless, the inclusion of anaerobic digestion could not substantially offset the energy demands related to aeration and pumping costs. This scenario was more similar to that with the MBR and WR2 processes than with WR1 (Figure 2, discussed next).

#### Membrane Bioreactor-based process (MBR): Greater Washington D.C. region, USA

Again, the two NSSS scenarios that resulted in the highest energy savings for the MBR process without optimization were gb-HRT and b-HRT (Figure 2, Table IV). On the other hand, the energy consumption due to nitrification could be reduced through urine separation, although not to the extent as achieved by influent load offsets through gb-HRT and b-HRT. Furthermore, implementation of internal recirculation alone did not have a substantial influence on energy demand, due to the influent load being conserved although with a reduction in influent flow rate (Figure 2). Combining urine-separation and internal recirculation, therefore, was less efficient than urine separation alone (Figure 2). The MBR unit process itself exerted a high energy demand, needed to maximize biomass retention. Therefore, overall, at this WWTP, even the inclusion of anaerobic digestion could not substantially offset the energy demands related to aeration and MBR operating costs (in contrast to an MBR-free process presented next for WR1), thereby leading to gb-HRT and b-HRT as the most effective NSSS options.

#### Water-Reuse 1 (WR1), California, USA

One of the highlights from the evaluation of upstream NSSS implementation at WR1 was the possibility to achieve <u>energy-positive centralized wastewater treatment</u> for the scenarios involving urine separation and internal recirculation coupled with urine separation (> 100% savings, Figure 2, Table IV). Such substantial energy efficiencies for these two scenarios were achieved due to the possibility of channeling away energy-intensive aerobic nitrogen conversion (nitrification), which is not needed for WR1 (unlike BNR1, MBR and BNR2), thereby allowing for the maximization of the beneficial impacts of biogas production and utilization. Therefore, in WWTPs with the possibility for biosolids conversion to biogas (or other energy endpoints) and where it might be possible to either offset the nitrogen load upstream using specific NSSS configurations, or where there are no limits on effluent nutrient loads, (for instance with WR1), the highest operating energy benefits could be achieved with RTs or NSSS that facilitate urine diversion and reuse. The ‘loss’ of such beneficial energy production is exemplified with the implementation of s-HRT at WR1, where the reduction in biogas production (owing to lower solids inputs) is reflected in higher energy requirements (negative savings, Figure 2). There was a low degree of improvement in the energy requirement associated with the implementation of b-HRT but not of gb-HRT (Figure 2). The observed changes were associated with (more) <u>proportionate</u> reductions in influent organic carbon, nitrogen, and phosphorus for gb-HRT. Finally, the impact of treatment and recirculation (without urine-separation) of part of the influent flow to the sewered WWTP also resulted in minimal changes to the energy requirements primarily owing to lack of a change in influent loads (Figure 2).

#### Water Reuse 2, (WR2): California, USA

Implementation of NSSS upstream did result in minimal energy savings (Figure 2). The two NSSS options that performed better than the rest were b-HRT, gb-HRT and s-HRT, although to a lower extent than that observed for the other WWTPs examined. (Figure 2, Table IV). Accordingly, the beneficial impact of upstream NSS implementation is expected to be minimal in such high-rate (low SRT) WWTPs that operate without the need for substantial organic carbon or nutrient conversion. On the other hand, the benefits of upstream NSS implementation even in such systems could be realized should these systems be required to transition or upgrade to more stringent treatment goals.

#### Multiplier Benefits of Integration are enable via WWTP Process Optimization

As seen in Table IV, higher degrees of NSSS implementation indeed resulted in higher absolute cost savings. On the other hand, through process optimization, the following observations were made that reflect the multiplier benefits of NSSS implementation coupled with optimization of the sewered WWTP. Essentially, with optimization, the degree of reduction in operating energy costs were higher than the corresponding degree of NSSS implementation (Figure 3). Furthermore, notwithstanding the higher absolute benefits at higher implementation, the relative benefits of optimization were highest at the lowest extent of NSSS implementation (Table IV, last column). Such results highlight the flexibility that NSSS implementation coupled with WWTP optimization provides and the need for a tailored solution. Decision-makers thus have the option between implementing broader coverage of NSSS or optimizing WWTP with lower NSSS coverage to attain similar cost benefits.

**Figure 3.**
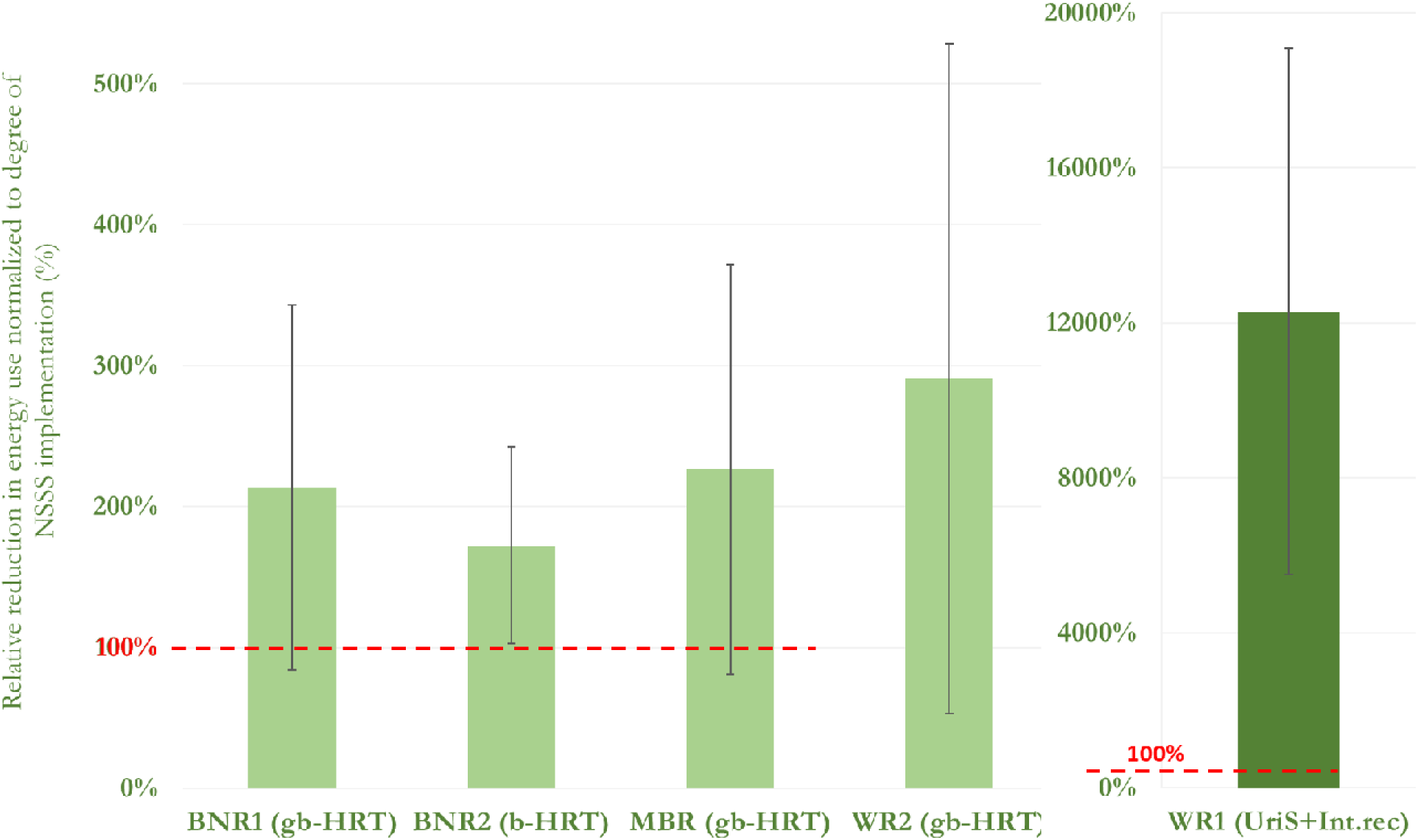
Relative savings in per capita energy costs normalized to degree of implementation (%) at the different sewered WWTPs tested as a function of different degrees of NSSS implementation <u>with optimization</u>. Results highlight the multiplier benefit of integrating NSSS and optimized WWTP operation (> 100%), with higher benefits at WR1. Height of the bars represents relative savings averaged over 10%, 30% and 50% the best non-optimized NSSS implementation. Error bars represent one standard deviation.

The benefits of optimization were the highest at WR2 (Figure 3, Table IV), essentially reflecting the fact that there is substantial opportunity to reduce energy demand at WR2 while continuing to achieve current performance or potentially even improving performance. Through optimization of the MBR-based WWTP, energy reductions were achieved by operation at lower MLSS concentrations in the MBR process and in the suspended growth reactors while meeting or exceeding the non-optimized WWTP performance.

The degree of improvement (compared to the non-optimized configuration) in energy demand for BNR2 upon optimization with gb-HRT and b-HRT was the lowest relative to the other WWTPs tested (Figure 3, Table IV). The main reason behind such limited improvement was the design configuration of the BNR2 process that resulted in kinetic limitation for nitrite oxidation with aggressive reductions in operating DO concentrations (data not shown). Nevertheless, optimization still resulted in operating cost savings through integration with b-HRT and gb-HRT (Figure 3, Table IV).

#### Overall perspectives

As observed from the results herein, across most centralized WWTPs configurations, improved wastewater treatment targets can be achieved by NSSS-mediated upstream treatment of sewage or both graywater and blackwater (gb-HRT) and just blackwater (b-HRT). On the other hand, urine separation by itself and when coupled with internal recirculation allowed for substantial savings in operating costs at WWTPs that focus on anaerobic digestion and don’t have effluent nutrient limits. The potential of s-HRT should be considered in the context of effluent discharge limits and energy recovery. Essentially, by channeling away some of the influent biodegradable solids and organic carbon, the efficiency of nutrient (both N&P) removal and energy recovery through anaerobic digestion could be compromised. Although not tested here, there could however be other operational benefits of s-HRT systems as related to conveyance and sewer maintenance costs.

Similarly, the main driver for internal recirculation could be to alleviate limitations in water availability for localized use rather than impacts on downstream WWTP alone. Treating and recycling the water at the point of NSSS-implementation when there is no change the influent *<u>load</u>* to the downstream WWTP should not result in a substantial impact as shown herein.

In addition to the operational savings demonstrated, coupling WWTP operations with upstream NSS implementation could offer increased flexibility in terms of adaptability to accept different outputs and options to achieve overall system process efficiency through novel system-integration approaches. This flexibility in operation could enable new, more beneficial, and holistic strategies for wastewater management, optimizing tradeoffs between different levels of centralized and decentralized resource recovery. For instance, one strategy could include targeting the NSSS to reduce flows (internal recirculation or urine and internal recirculation) which could enable to focus the WWTP process on large scale resource recovery from high load influent, rather than on nutrient removal. Such a strategy could support energy-recovery at WWTPs coupled with nutrient recovery to increase economic benefits in tandem. An interesting possibility for such integrated NSSS-WWTPs could be efficient co-treatment of internal recycle flows (such as digestate, filtrate or centrate) or even high strength agricultural influents or food- and animal-waste as applicable.

Another possibility for integrated NSSS-WWTPs could also be the prospect of improved climate-change adaptation via stormwater flow management at the centralized WWTPs using the excess capacity made available through upstream treatment capacity offsets. Alternatively, upon integration, the downstream WWTP processes could also be oriented towards industrial or potable water reuse. Such a strategy might be preferable for rural areas with proximal agricultural practice.

In summary, this study demonstrated the beneficial impacts relating to both operational flexibility and cost savings of integrating of NSSS with sewered WWTPs. NSSS-WWTP integration provides water and wastewater utilities additional options to ensure access to wastewater treatment and sanitation especially in light of pressures including increased service demand, associated with expanding populations, more stringent treatment limits, potentially exacerbated by climate-variability or resource-limitations. In turn the operational savings realized at sewered WWTPs can help incentivize the broader adoption and implementation of NSSS, thereby fostering a more resilient wastewater and sanitation service and infrastructure.

## Methods

The impact of six NSSS technologies (Table I) was examined on the performance of each of five centralized WWTP configurations that represent a broad spectrum of the most common biological wastewater treatment processes sector-wide (Grady et al. 1999, Tchobanoglous et al. 2003) (Table I). Six NSSS adoption scenarios were tested based on their ability to independently treat grey- or black-water, solids, or urine, with or without internal greywater recycling (Table II). All the NSSS examined entailed target liquid and solid treatments allowing for safe discharge, recycling, or reuse onsite. For each centralized WWTP-NSSS combination, three degrees of NSSS implementation of 10%, 30%, and 50% were evaluated. For each NSSS, reductions in the influent flow and concentrations to the sewered WWTP were calculated based on the chosen degrees of NSSS implementation (Tables II-III).

As a first step, process models of each WWTP were developed based on influent flows and loads, process configurations, and process performance, all acquired directly from the WWTPs themselves (workflow described in Figure 1). The principal state-variables tracked included concentrations of COD, total Kjeldahl nitrogen (TKN), total phosphorus (TP), mixed liquor suspended solids (MLSS), including total, volatile and inert fractions. The biological transformations in the process simulations were captured based on a modified version of the Activated Sludge model (Henze et al. 1987).

### Details of the centralized WWTPs evaluated in this study

#### 1. BNR1, Beijing, China

The overall process configuration at this WWTP (labeled BNR1) is a variant of a four-stage Bardenpho process and includes, pre-denitrification, nitrification and aerobic organic, carbon oxidation, post-denitrification and re-aeration bioreactors. The WWTP serves a calculated population size of 289,000 individuals while processing an average flow rate of 60 million liters per day (ML/d), containing a nitrogen load of 2700 kg-N/d. This facility represents a case when the influent COD:N is limited (COD:TKN=5:1), and nitrogen removal is required by regulation.

#### 2. BNR2: Durban, South Africa

This WWTP (BNR2) employs a variant of the Modified Ludzack-Ettinger (MLE)-based BNR process, serving roughly 328,000 population equivalents. Biosolids management entails anaerobic digestion. A portion of the biogas produced in the primary digesters is used for heating the digesters while the excess is flared. All internally produced streams such as thickener overflows and post-digestion-thickener overflows are channeled to primary settling tanks. The secondary effluent is discharged to the series of three maturation ponds and the pond effluent is chlorinated prior to discharge to the Umhlanga River in Durban, SA.

#### 3. MBR-based process, Greater Washington, D.C. region, USA

This WWTP (labeled MBR) is based on an energy-intensive Membrane Bioreactor process and serves a calculated population size of 195,000 individuals, treating an average flow rate of 30 ML/day, containing a nitrogen load of 1828 kg-N/d. The treatment process is comprised of bioreactors geared towards BNR operations coupled with a membrane bioreactor for efficient solids-liquids separation. Primary and secondary solids are anaerobically digested for energy recovery.

#### 4. WR1: California, USA

This WWTP represents an advanced activated sludge treatment plant for direct potable reuse, labeled (WR1) serving a calculated population size of 2.7 million individuals, treating an average flow rate of 454 ML/day, containing a nitrogen load of 25,435 kg N/d. The liquid-phase biological treatment component of WR1 is comprised of a trickling filter and suspended phase aerobic and anoxic processes. The plant is designed to achieve some degree of nitrification as well as BNR. Primary and secondary solids are anaerobically digested for energy recovery.

#### 5. WR2: California, USA

The second water reuse facility labeled WR2 employs a high-purity oxygen activated sludge (HPOAS) to convert ammonia to nitrate, and serves a calculated population size of 1.3 million individuals, treating an average flow rate of 223 ML/day, containing a nitrogen load of 130,850 kg-N/d. For this WWTP, the use of a high-purity oxygen activated sludge (HPOAS) is a distinct feature. Primary and secondary solids are anaerobically digested for energy recovery.

### Model Calibration

For each WWTP, a process model was generated and calibrated, as described previously for a NSS process converting fecal sludge to volatile fatty acids in Kumasi, Ghana(Shih et al. 2017). Briefly, calibration for each WWTP model entailed comparing the measured influent characteristics and process performance (provided by the WWTPs) with the simulated process performance and modifying the WWTP model parameters <u>as minimally as possible</u> to achieve the best correspondence between measured and simulated data. Process performance measures that were compared and reconciled included concentrations of effluent organic carbon (expressed as chemical oxygen demand, COD), ammonia-N, nitrite-N, nitrate-N, total nitrogen, total phosphorus, total suspended solids; total and volatile suspended solids concentrations in the activated sludge bioreactors, and total and volatile solids, total gas production and gas composition in anaerobic digestion. Based on the calibrated model, we additionally calculated and tracked operating energy costs as a function of NSSS integration.

### Determining the impact of different NSSS on WWTP operating costs and performance

Subsequently, the calibrated model was used to assess the effects of different extents of NSSS implementation on WWTP performance. For comparison among the scenarios, we largely maintained a constant solids residence time (SRT) within each WWTP process. For the two NSSS that include urine separation, documented urine characteristics were used (Table III).

### Optimization of centralized WWTP operation to maximize operational efficiency

Further process optimization was conducted to quantify enhanced operational efficiency and cost savings associated with WWTP operations when integrated with different NSSS options. As part of optimization, we explored changes in process operations, based on existing infrastructure, that would enable the WWTPs to achieve similar non-optimized performance with lower energy usage for the top two most effective NSSS options.

## Acknowledgements

Support for this work by the Bill and Melinda Gates Foundation Grant INV 035862 is gratefully acknowledged.

